# Network analyses implicate a role for PHYTOCHROME-mediated light signaling in the regulation of cuticle development in plant leaves

**DOI:** 10.1101/812107

**Authors:** Pengfei Qiao, Richard Bourgault, Marc Mohammadi, Laurie G. Smith, Michael A. Gore, Isabel Molina, Michael J. Scanlon

## Abstract

Plant cuticles are composed of wax and cutin, and evolved in the land plants as a hydrophobic boundary that reduces water loss from the plant epidermis. The expanding maize adult leaf displays a dynamic, proximodistal gradient of cuticle development, from the leaf base to the tip. Laser microdissection RNA Sequencing (LM-RNAseq) was performed along this proximodistal gradient, and complementary network analyses identified potential regulators of cuticle biosynthesis and deposition. Correlations between cuticle development and cell wall biosynthesis processes were identified, as well as evidence of roles for auxin and brassinosteroids. In addition, our network analyses suggested a previously undescribed function for PHYTOCHROME-mediated light signaling during cuticular wax deposition. Genetic analyses reveal that the *phyB1 phyB2* double mutant of maize exhibits abnormal cuticle composition, supporting predictions of our coexpression analyses. Reverse genetic analyses also show that *phy* mutants of the moss *Physcomitrella patens* exhibit abnormal cuticle composition, suggesting a role for light-stimulated development of cuticular waxes during plant evolution.

## Introduction

Light perception plays important roles in the regulation of plant metabolism and development (1–5), including the activation of lipid production in algae (6, 7). However, aquatic algae lack cuticles, the hydrophobic barrier deposited on the epidermis of all land plants that prevents nonstomatal water evaporation through the plant surface. Cuticles cover the above ground shoot of land plants, and enabled their invasion and colonization of the dry and hostile terrestrial environment. Since the majority of water loss in plants occurs through the epidermis, the cuticle imparted a significant advantage in plant evolution by providing a barrier to desiccation (8–12).

The plant cuticle comprises a mixture of solvent-soluble lipids (waxes) plus a lipid polymer (cutin). Waxes are long-chain, non-polar molecules, composed mainly of hydrocarbons (alkanes and alkenes), aldehydes, alcohols, ketones and wax esters. In contrast, cutins are polymers of hydroxy fatty acids connected by ester bonds (13–15). Waxes and cutins are both formed *de novo* from long-chain C16 and C18 fatty acids synthesized within plastids of the plant epidermis (16, 17). In *Arabidopsis thaliana*, these long-chain fatty acids are converted to CoA thioesters by LONG-CHAIN ACYL-COENZYME A SYNTHASE (LACS), and subsequently transported into the endoplasmic reticulum, where they are elongated by the fatty acid elongase (FAE) complex (18). After elongation and further modification, these cuticle lipids are exported out of the plasma membrane and into the apoplastic space, where small protein transporters such as LTPs may facilitate the transport of lipid precursors to the site of cuticle deposition (19–21). Cuticle development is regulated by many factors such as the phytohormone abscisic acid (ABA), water deficit, osmotic stress, and light (18, 22, 23).

Previous studies in Arabidopsis have shown that the expression of several cuticle biosynthesis genes is induced by light (18, 22, 23). Intriguingly, light-activated photoenzymes can stimulate the enzymatic conversion of fatty acids to hydrocarbons in green algal relatives of land plants (6, 7), although these lipids are not deposited on algal surfaces to form a cuticle. Likewise, PHYTOCHROME light receptors, which regulate a variety of physiological processes during plant growth and development, are also found in green algae (24, 25). LTPs, on the other hand, are only found in land plants, and are proposed to play a pivotal role in cuticle biosynthesis (26–30).

In this study, we utilized the expanding adult leaf as a model system to elucidate the spatial-temporal gradient of maize cuticle development. Transcriptomic analyses were performed along the proximodistal axis of the developing maize leaf eight as it emerged from darkness to light. Complementary network analyses of transcriptomic data were employed to identify patterns of epidermal gene expression underlying the cuticle composition gradient previously identified in this tissue (31), and to identify novel candidate genes for cuticle development in maize. Network analyses suggested a previously unidentified role for PHYTOCHROME during cuticle development, which was confirmed by genetic and biochemical investigations in the evolutionarily divergent model plants *Zea mays* and *Physcomitrella patens*. We propose a mechanistic model whereby phytochrome-mediated light signaling in land plants was a critical step enabling the evolution of cuticle development.

## Results and Discussion

### Transcriptomic analyses of cuticle development in the adult maize leaf

Previous analyses demonstrated a gradient of cuticle composition along the proximodistal axis of the expanding leaf eight of maize inbred line B73, from light-shielded proximal intervals to light-exposed distal regions (Figure 1A). In general, longer-chain wax components and cutin monomers increase in abundance as the leaf transitions from the dark to light (31). To capture the transcriptional gradient coinciding with these biochemical changes in cuticle composition, seven developmental stages along the proximodistal axis of leaf eight were laser-microdissected for RNAseq analysis. Each stage comprised a two-centimeter-long interval, collected between 2 and 22 cm from the leaf base, encompassing the point of emergence of leaf tissue into the light at ~18 cm (Figure 1A). For each of the seven proximodistal intervals examined, an L1-derived epidermal sample and an L2-derived internal sample were microdissected (*SI Appendix*, Fig. S1), followed by RNA sequencing to construct their transcriptomes. Principal component analysis (PCA) identified two PCs that collectively explain 60.29% of the total sample variance in the transcriptomic data. Specifically, the first PC explains 38.22% of the total sample variation, and corresponds to the seven proximodistal leaf intervals analyzed, whereas the second PC (PC2, 22.07% of sample variance) delineates epidermal and internal tissues for each leaf developmental stage (*SI Appendix*, Fig. S2). These data show that in addition to the biochemical gradient in cuticle composition, the leaf intervals examined in maize leaf 8 also exhibit a transcriptomic gradient.

**Fig. 1.**
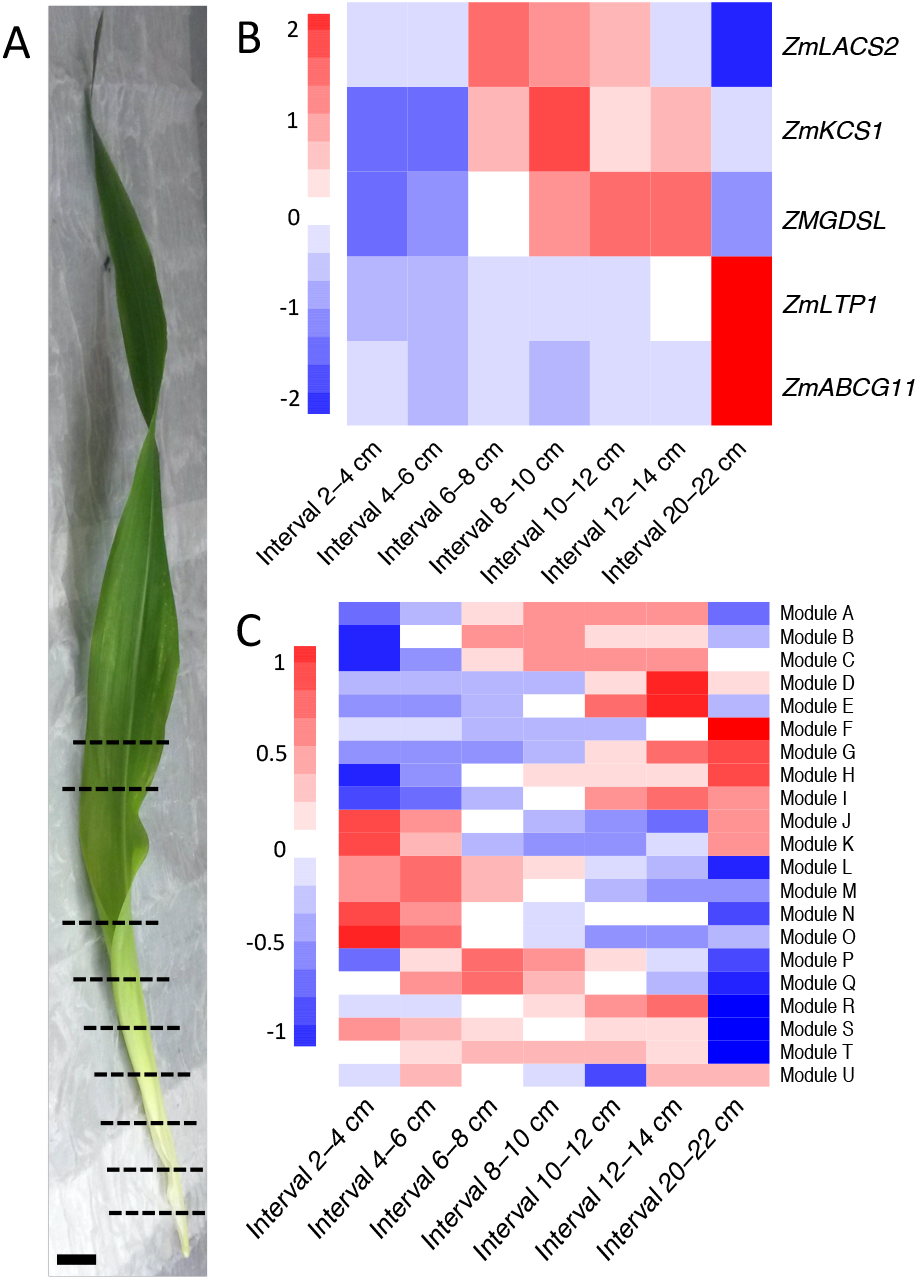
Proximodistal transcriptomic gradients in the expanding adult maize leaf. (A) Dashed lines demarcate the boundaries of 2-cm intervals collected along the proximodistal axis of the expanding maize leaf eight (note: the leaf emerges from the whorl into the light ~18 cm from the leaf base). Scale bar (solid line) = 2 cm. (B) Heatmap showing the expression patterns of selected cuticle biosynthesis and transport genes across seven proximodistal intervals of leaf 8. (C) Heatmap showing the expression levels of eigengenes for the 21 modules identified in our gene co-expression network.

### Identification of putative cuticle regulatory genes across the leaf developmental gradient via directed network inference

A gene regulatory network (GRN) was constructed using causal structure inference (CSI), as a means to identify candidate regulators of cuticle biogenesis based on analysis of the epidermal transcriptomic patterns in the seven intervals of maize leaf eight. The CSI algorithm accounts for the biological delay of one gene’s effect on another gene’s expression level, by incorporating both the spatial and temporal transcriptomic information within all the sampled leaf intervals (32). CSI generates a network with connections (edges) between genes (nodes), directed from putative regulators toward their targets. A Bonferroni-corrected significance level was used to identify putative regulatory relationships among epidermally-enriched transcripts from the seven leaf intervals, since the epidermis is the site of cuticle biosynthesis (16, 33).

Previously described cuticle biosynthetic genes were differentially expressed across the seven leaf intervals (Fig. 1B), coinciding with the changes in cuticle composition in the expanding leaf eight. For example, expression levels of ABCG transporters (e.g. *ZmABCG11* in Fig. 1B) that deliver wax components and cutin monomers out of the plasma membrane (34–36) show a consistent increase that correlates with the accumulation of cutin monomers (31) within these intervals. These data established that there is a gradient in the epidermal gene expression of leaf 8, especially for genes involved in cuticle biosynthesis and deposition. The gene expression gradients can be incorporated into the GRN model as the “biological delay” of their regulators’ functions, supporting the efficacy of our CSI approach to utilize the regulatory delay. Known cuticle biosynthetic genes were used as “baits” to identify putative regulators of cuticle biogenesis. Baits included *ECERIFERUM* (37) (GRMZM2G029912) and *BETA-KETOACLE REDUCTASE* (*KCR*) (38) (GRMZM2G090733), and are highlighted in yellow in Figure 2. Known cuticle biosynthesis nodes showed incoming edges from genes with no previously described function in cuticle development (such as GRMZM2G055469, GRMZM2G120619, GRMZM2G078959), identifying these new genes as potential regulators of cuticle biosynthesis. Although many of these candidate genes are not previously described to function during the regulation of cuticle development, others such as GRMZM2G055158, have homologs in Arabidopsis (*MYB20*) that are involved in secondary cell wall thickening (39). Thus, these genes and several others with numerous outgoing edges (i.e. hubs, *SI Appendix*, Table S1), represent potential candidate genes for reverse genetic analyses of maize cuticle biosynthesis.

**Fig. 2.**
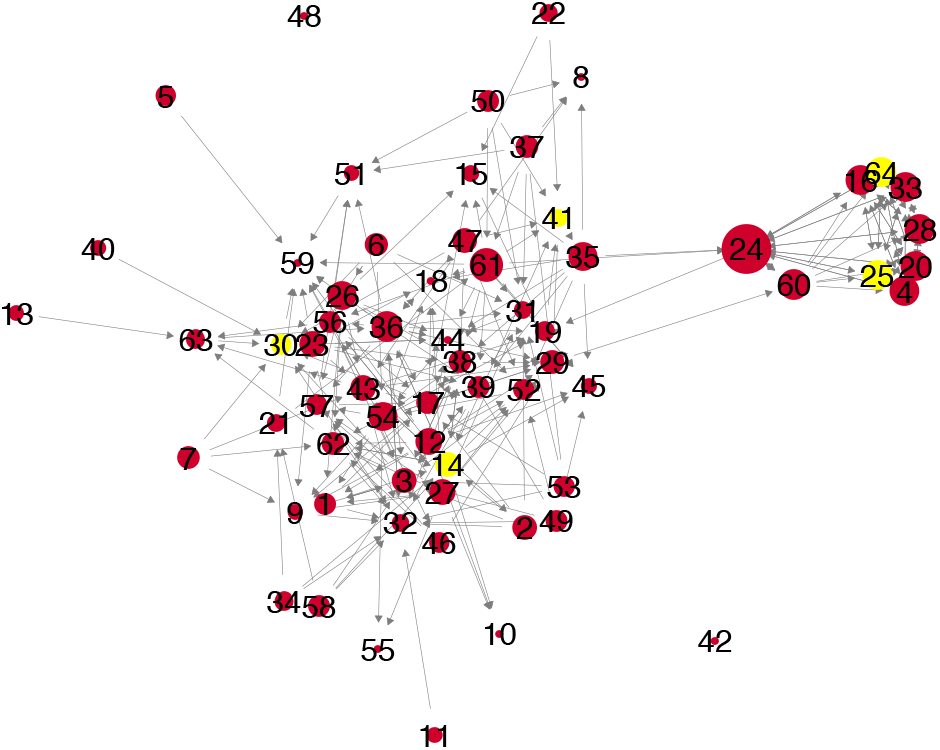
Gene regulatory network of epidermally enriched genes in the maize expanding adult leaf 8. Previously described genes functioning in cuticle biosynthesis or lipid transport, and cuticle regulatory genes are highlighted in yellow. Arrows point to potential targets of the corresponding regulatory genes. The size of the nodes correlates with the number of targets for each node (directional edges coming out of the node), i.e. outdegree. Each number corresponds to a specific gene identified in *SI Appendix* Table S1 (numbers on nodes correspond to numbers in column A of Table S1).

### A weighted co-expression network analysis identifies additional candidate regulatory genes for cuticle biosynthesis

Although the CSI algorithm is a powerful tool enabling gene discovery and their functional relationships, its ability to model the biological delay of a gene’s effect can also be a disadvantage, depending upon how the transcriptomic samples are collected. For example, although CSI will detect the delayed effects of a gene *between* two 2 cm leaf intervals, it will fail to detect these effects *within* any 2 cm leaf sample. Hence, we utilized a second, complementary method called gene co-expression network (GCN) analysis (40, 41) to identify additional candidate genes involved in regulating the biosynthesis of the maize cuticle. GCN is essentially a “guilt-by-association” approach, wherein correlations in gene expression levels implicate co-regulation of gene pairs within the network.

We constructed a weighted gene co-expression network based upon the expression-level correlations of all 11,816 epidermally-transcribed genes identified in our LM-RNAseq (40). The transcriptome was partitioned into 21 modules (*SI Appendix*, Table S2 - 22); Figure 1C illustrates the expression levels of eigengenes (idealized representative genes) within these 21 modules at each of the seven, leaf developmental stages analyzed. Comparisons of gene expression levels to cuticle lipid profiles at each interval reveals interesting correlations, as shown in Figure 3. For example, expression levels of genes in modules C, H and I are positively correlated with the accumulation of wax esters, whereas modules L, M, N and O show the opposite trend. Many modules with positive or negative correlations with specific cuticle lipid profiles contain known cuticle biosynthesis and regulatory genes, for example, in module H, *KCS* homologs GRMZM2G104626 and GRMZM5G894016 show a correlation coefficient of 0.94 and 0.69 with the cutin monomer hydroxy fatty acid 16:0 16-OH. The strategic use of this network analysis for gene discovery is described below.

**Fig. 3.**
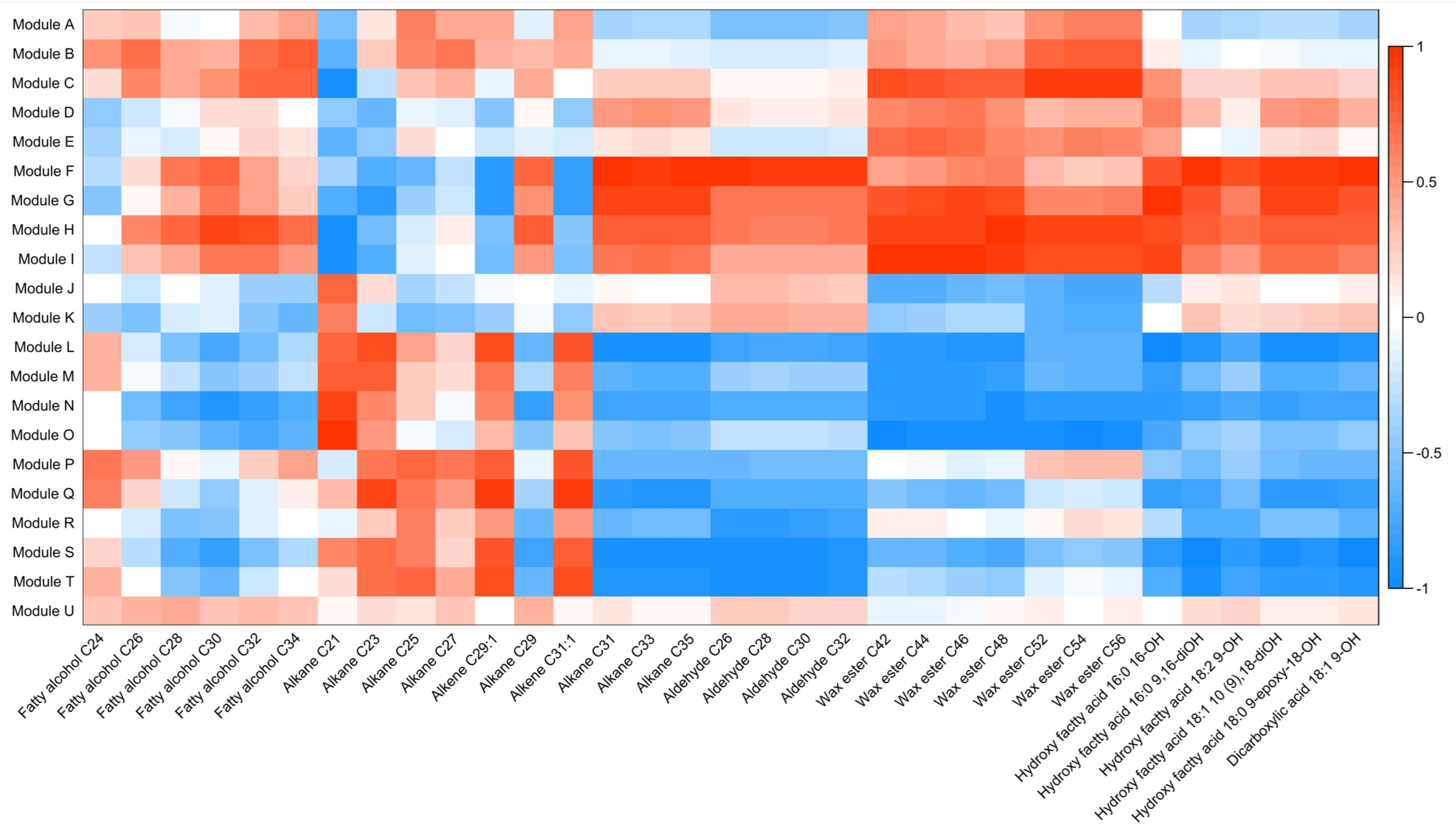
Heat map depicting the correlation of each cuticle lipid component (x axis) to the 21 co-expression modules (y axis) identified in transcriptomic analyses of the expanding maize leaf 8. Colors (red to blue) correspond to the values of the Pearson’s pairwise correlations, where red (+1) is positively correlated and blue (−1) is negatively correlated. Cuticle lipid abundance data used for this analysis are in (31).

One such strategy is to interrogate the direct neighbors of known cuticle biosynthetic genes within the network, as demonstrated in analyses of five modules (Q, F, C, A and I) showing enrichment for known cuticle genes. For instance, members of several gene families known to function in cuticle biosynthesis are overrepresented in Module F. These include *KCS*s, *ABC TRANSPORTER*s, and *LTP*s (18, 19, 36), whose direct neighbors exhibit strong co-expression with these cuticle genes, indicating regulatory roles or co-regulation. The direct neighbors of known cuticle biosynthesis genes in these modules are summarized in *SI Appendix*, Table S23, and comprise additional potential candidate genes involved in the regulation of cuticle biosynthesis in the expanding maize leaf (40, 42).

A second gene discovery approach enabled by GCN analyses is to examine the hubs within the network, defined as the most connected nodes that are essential to network function (43, 44). In our case, the hubs of modules that are significantly correlated with cuticle components are usually a subset of the candidates identified via the “direct neighbor” approach described above.

For example, in module F many of the hubs are also direct neighbors of *KCS*s, *ABC TRANSPORTER*s, and *LTP*s. These findings further confirm the importance of module F, and especially its hubs, on cuticle development in the expanding maize leaf. A summary of candidate genes for cuticle development for all the modules with significant correlation with cuticle components is presented in *SI Appendix*, Table S2. Two examples of GCN-based identification of candidate cuticle synthesis and regulatory genes are discussed below.

Cuticles are deposited on epidermal cell walls, and thus can be regarded as plant cell wall modifications. Our data suggest that cuticle development and cell wall biosynthesis may indeed be coupled. Six SAUR genes, a gene family that is induced by auxin and implicated in cell wall expansion (45), share direct neighbors with *KCS*s and *MYB* transcription factor genes (*ZmMYB96*, GRMZM2G098179 and GRMZM2G139284) with homology to Arabidopsis regulators of wax biosynthesis (46). Previous studies showed that WAX INDUCER1/SHINE1 transcription factors regulate cuticle biosynthesis, and also induce the expression of several genes encoding pectin-modifying enzymes (15). Intriguingly, two maize pectin lyase-like superfamily protein encoding genes occupy central positions in our GCN, as do genes controlling synthesis of the cell wall constituents xyloglucan and cellulose (*SI Appendix*, Table S2).

Our GCN also suggests a role for the phytohormones auxin and brassinosteroid (47) during cuticle biosynthesis. AUX/IAAs are essential transcriptional repressors during auxin signaling, and are degraded in the presence of auxin (48). Two *AUX-IAA* transcripts (GRMZM2G159285, GRMZM2G077356) are identified as key hubs and direct neighbors in our GCN, and comprise excellent candidate genes for cuticle development. Likewise, the central positions of *BRASSINOSTEROID-RESPONSE RING-H* homologs in our GCN suggest a previously undescribed role for brassinosteroids during cuticle biosynthesis.

### Light-regulated cuticle development: *phytochrome* (*phy*) mutants have altered cuticle composition

Previous studies showed that light induces cuticular wax biosynthesis in land plants, and the expression levels of several fatty acid elongase complex transcripts decrease in dark-grown plants, thus reducing the amount of cuticular wax (18, 22, 23, 49). Moreover, biochemical analyses also revealed that longer chain wax components are more abundant in the distal, light-exposed intervals of maize leaf 8 (Fig. 1A) (31). Although algal relatives of the land plants do not develop a cuticle, light exposure does induce the conversion of hydrocarbons from long-chain fatty acids (6, 7). A GO term analysis of module H showed significant enrichment of transcripts that respond to light stimuli (*SI Appendix*, Fig. S3), and our GCN identified a previously-undescribed correlation between *PHYTOCHROMEB1* and cuticle development (*SI Appendix*, Table S24). PHYTOCHROMES (PHYs) are red/far-red light photo-reversible chromoproteins that regulate gene expression in response to light. Maize contains six *PHY* homologs (*PHYA1*, *PHYA2, PHYB1, PHYB2, PHYC1, PHYC2*) (50), and *PHYB1* occupies a central position in module H (Fig. 4A). Module H is also strongly correlated with many cuticle components, such as C_31_ - C_35_ alkanes (Fig. 3). Gas chromatography–mass spectrometry (GC-MS) comparisons were performed on flag leaves of the previously described *phyB1 phyB2* double mutant and non-mutant sibling flag leaves (50). The *phyB* double mutant cuticles showed significant differences at 5% FDR in several wax components. Lower amounts of alkanes and fatty alcohols were observed for species with 32 carbons and longer; only the C_34_ aldehydes were reduced whereas the wax esters were unaffected. In addition, a likely compensatory increase in shorter-chain components was observed, including: C_24_ - C_30_ fatty alcohols (Fig. 4B), C_23_ - C_31_ alkanes, C_28_ - C_32_ aldehydes, C_41_, C_43_, C_45_, C_47_, C_48_, and C_50_ wax esters (*SI Appendix*, Fig. S4-6) when compared with sibling leaves. These data strongly support models wherein light regulates cuticle biosynthesis, as predicted by module H.

**Fig. 4.**
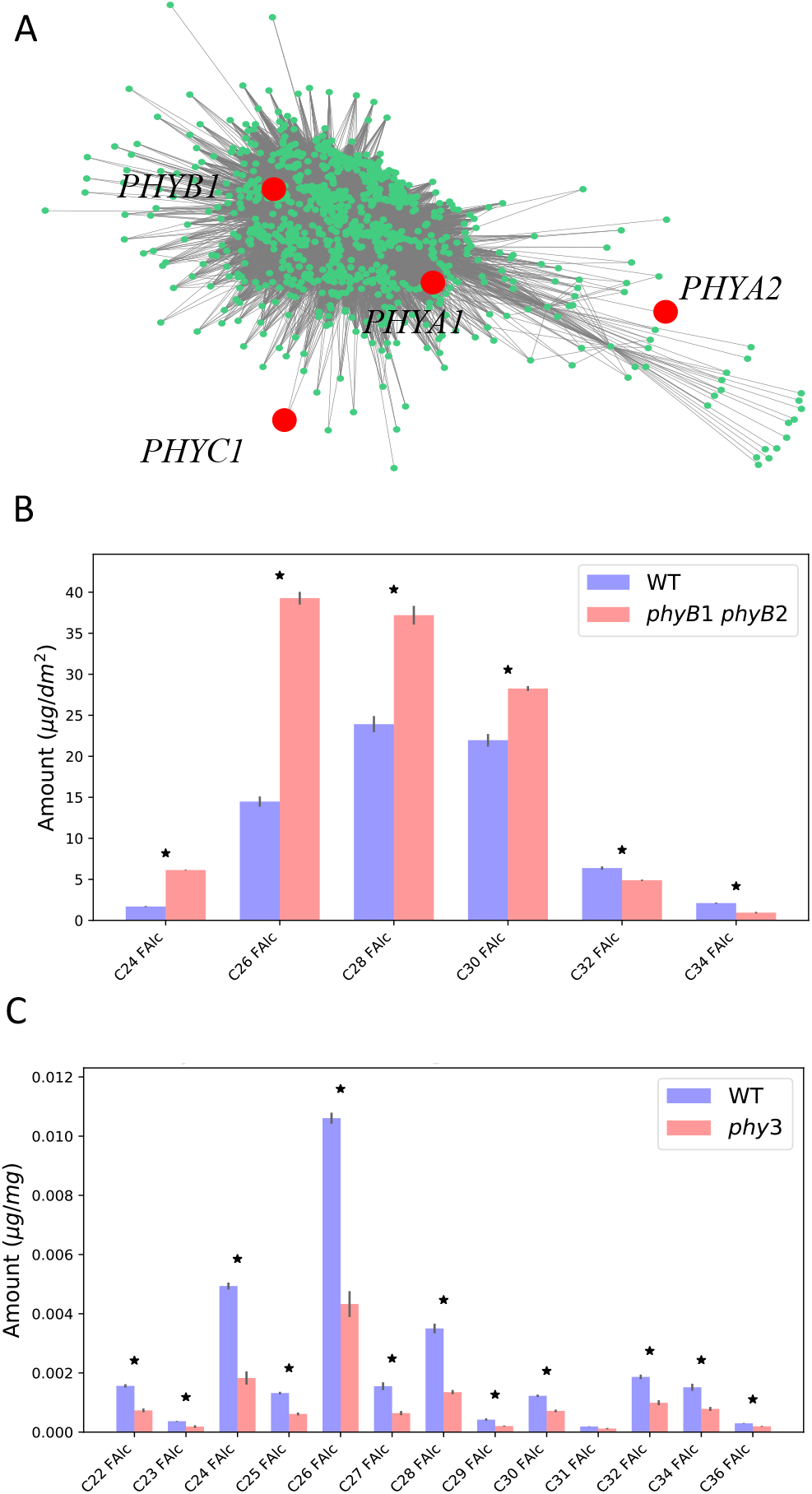
Phytochrome regulates cuticle wax composition in the adult maize leaf 8. (A) Visualization of the co-expression network of module H. The red colored nodes correspond to the phytochrome homologs *PHYA1, PHYA2, PHYB1*, and *PHYC1*, some of which (*PHYB1* and *PHYA1*) occupy central positions of the network (with numerous connections with other nodes). Module H is also significantly correlated with specific cuticle components (Fig. 3). (B) Comparisons of the fatty alcohol cuticular wax profiles in leaves of *phyB1 phyB2* double mutant versus nonmutant maize. (C) Comparisons of the fatty alcohols in cuticular wax profiles in gametophyte colonies of *phy3* mutant versus non-mutant *Physcomitrella patens*. Error bars represent standard errors. Asterisks indicate significant differences (< 0.05 FDR) between samples in unpaired t-tests.

Interestingly, the defects in cuticle composition observed in the *phyB1 phyB2* double mutant mirror the changes in cuticle components as the leaf emerges from the whorl. That is, the amounts of longer-chain wax components increase as the leaf is exposed to light (31), while the *phyB* double mutant shows defects in wax components that require fatty acid chain elongation beyond 32 carbons.

To determine whether PHY regulation of cuticle accumulation is simply a maize-specific phenomenon or is in fact found in other land plants, equivalent analyses of cuticle lipids were performed on *phy3* mutant colonies of the moss *Physcomitrella patens* (2), a member of the bryophytes that diverged from later-evolved plant lineages early in the evolution of land plants (29). Intriguingly, *phy3* mutants showed significant reductions at 5% FDR in the amounts of all but one of the identified fatty alcohols (Fig. 4C), including: C_29_, C_31_, and C_33_ alkanes, C_25_, C_26_, C_27,_ C_28_ and C_30_ aldehydes, and C_38_, C_40_, C_41_, C_42_, C_43_, C_44_, and C_45_ wax esters (*SI Appendix*, Fig. S7-9) when compared with wild-type plants. These *phy3* mutant moss phenotypes suggest that PHY regulation of cuticle development may be a widespread phenomenon of land plants, although analyses of additional plant taxa are required to rigorously test this model.

PHYTOCHROMES are found in all green plants including algae (24, 25), although cuticles and are an evolutionary innovation of land plants (9, 11, 26–30). Although light signaling-associated changes in lipid biosynthesis are present in the green algae ancestors of land plants, these light-induced lipids are not utilized to form a cuticle in algae. Our data from bryophyte mosses and angiosperm grasses suggest that cuticle development in land plants is regulated by PHYTOCHROME-mediated light signaling, as an innovation during the evolution of land plants.

## Materials and Methods

### Plant material and growth conditions

B73 seeds were obtained and from the Maize Genetics Cooperation Stock Center; maize plants were grown in a 25 °C day, 20 °C night, in 60% relative humidity and 10-hour day length Percival A100 growth chambers (Percival Scientific, Perry, IA) until harvest.

Maize *phyB1 phyB2* mutants and their non-mutant siblings were obtained from J. Strable and T. Brutnell (50), moss *phy3* mutants and their non-mutant siblings were obtained from J. Hughes and grown as described (51).

### Laser microdissection and RNAseq analysis

The unexpanded eighth leaf of the inbred B73 maize plant was harvested in three biological replicates, three plants per replicate, when the leaf was 45~55 cm, around 33 days after planting. Leaf 8 was dissected out of the whorl and segmented into 2 cm long intervals, up to 22 cm from the leaf base, seven of which (six intervals from 2 - 14 cm, and one interval from 20 – 22 cm) were fixed and paraplast–embedded for use in laser microdissection as described (52). Epidermal and internal tissues were microdissected; RNA was extracted using the PicoPure™ RNA isolation kit and linearly-amplified using the Arcturus RiboAmp ® HS PLUS RNA Amplification kit. RNA Sequencing libraries were constructed with the NEBNext Ultra™ RNA Library Prep Kit for Illumina, and the HiSEQ 2500 instrument was used for sequencing. RNAseq reads were deposited in the NCBI SRA under accession number SRP116320 (https://www.ncbi.nlm.nih.gov/Traces/study/?acc=SRP116320&o=acc_s%3Aa).

### Wax extraction and analysis

Waxes were extracted by submerging the tissue in pure chloroform for 60 seconds, followed by evaporation under a gentle stream of nitrogen. Dry wax samples were analyzed with GC-MS as described previously (31).

### Differential expression analysis

Differential expression analysis was performed with edgeR 3.3.2 package in R (56, 57). Genes with counts fewer than 2 counts per million reads (cpm) were filtered out and analysis was carried out under FDR < 0.1 as the significant measure.

### Weighted co-expression network analysis

The correlation between genes was performed using a modified version of Tukey’s Biweight correlation (58), which was later used to calculate the distance matrix. The calculations were done using WGCNA 3.3.0 package in R (40, 41). The distance matrix was used for the dynamic hierarchical clustering and construct the edges (connections) between nodes (genes) in the network. Network analysis of hubs and direct neighbors was done in Python 2.7 using NetworkX 1.11 module (59).

### Casual structure inference analysis

The epidermally upregulated genes were enriched with differential expression analysis comparing epidermal and internal samples, with Bonferroni-correction. For pair-wise comparison between adjacent epidermal sections, genes that showed significant differences in at least one comparison were input into the CSI algorithm using Cyverse (32). Each pair of genes was taken as a prior, and their effects on the section-derivative of the expression levels of all other genes gene was modeled using a Bayesian non-parametric approach (32). Each gene’s marginal effect on another genes expression gradient was calculated, and treated as a regulatory edge in the GRN.

## ACKNOWLEDGMENTS

The work was funded by NSF IOS award #1444507. We thank all members on the cuticle project for discussion and inputs, especially S. Matschi, M. Vasquez, A. Nguyen, M. Lin. We thank S. Leiboff for help in RNAseq processing. We also thank J. Strable, T. Brutnell for the *phyB1 phyB2* double mutant and sibling maize seeds, J. Hughes for the *phy3* moss mutant, and J. Cammarata for the help of growing moss.

