## Supplemental Figures 1-2, 4-9 for "Network analyses implicate a role for PHYTOCHROME-mediated light signaling in the regulation of cuticle development in plant leaves"

**This PDF file includes:**

Figs. S1 to S2, S4 to S9

**Other supplementary materials for this manuscript include the following:**

Fig. S3
Tables S1 to S24

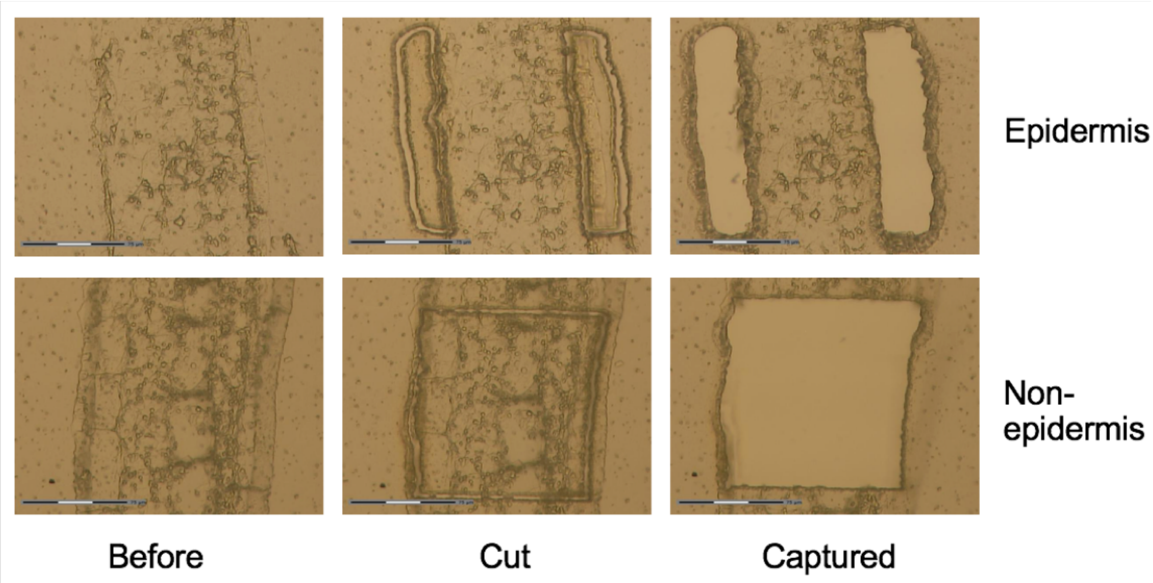


**Fig. S1.** Laser microdissection was performed on leaf tissues to isolate the L1-derived epidermal layers (top row) and L2-derived internal layers of each targeted leaf interval along the proximodistal axis of the expanding leaf 8. From left to right, each column corresponds to a leaf section before, during and after microdissection. Scale bars = 75 µm.

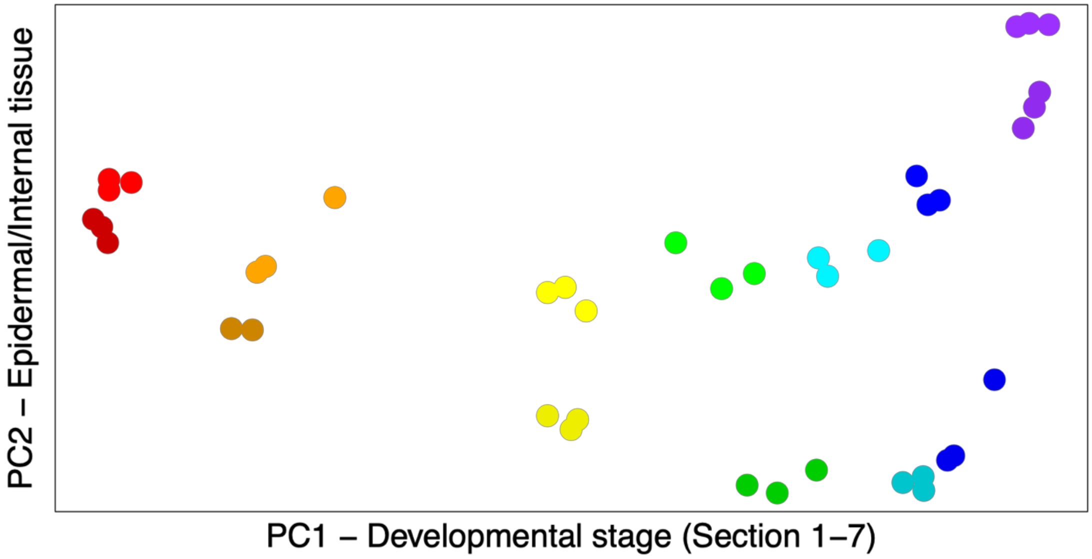


**Fig. S2.** Principal component analysis identified two principal components (PCs) corresponding to the developmental stage (PC1) and tissue type (PC2) of leaf 8 samples in our LM-RNAseq analysis. Each point corresponds to one RNAseq sample. From left to right, each color corresponds to a specific developmental stage (youngest to oldest part of the leaf). From bottom to top, darker color shades represent L1-derived epidermal tissues, and lighter shades represent L2-derived internal tissues. For interval two, one outlier epidermal sample was removed.

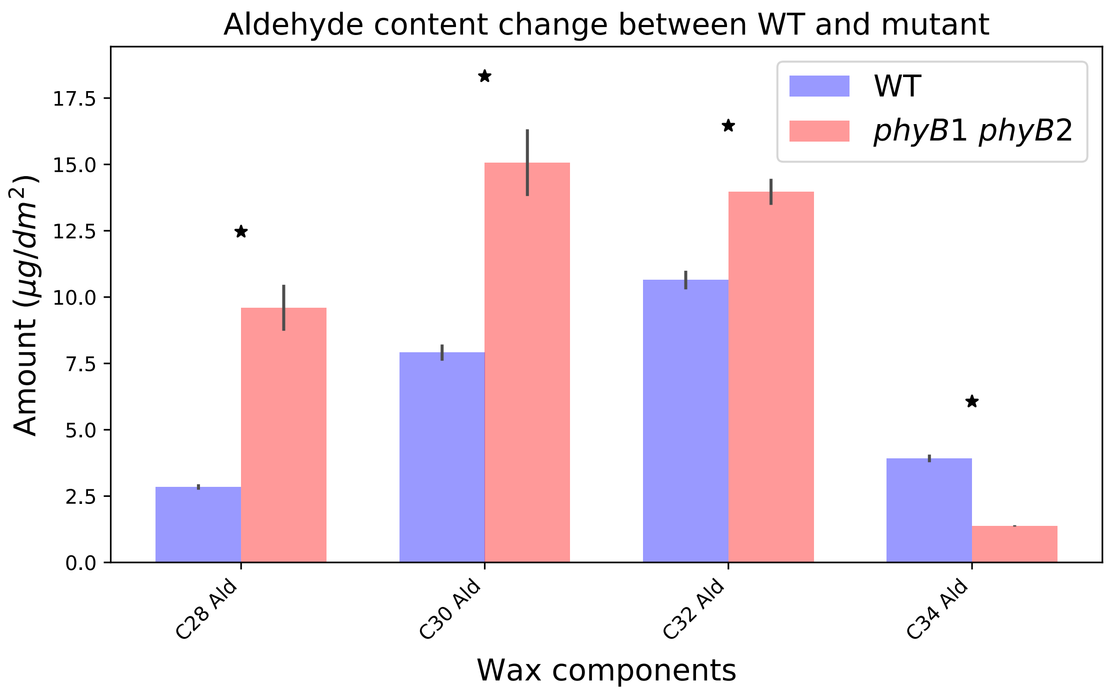


**Fig. S4.** Comparisons of aldehydes in cuticular wax profiles in *phyB1 phyB2* double mutants versus nonmutant plants in maize. Asterisks indicate significant differences (< 0.05 FDR) between samples. Error bars represent standard errors.

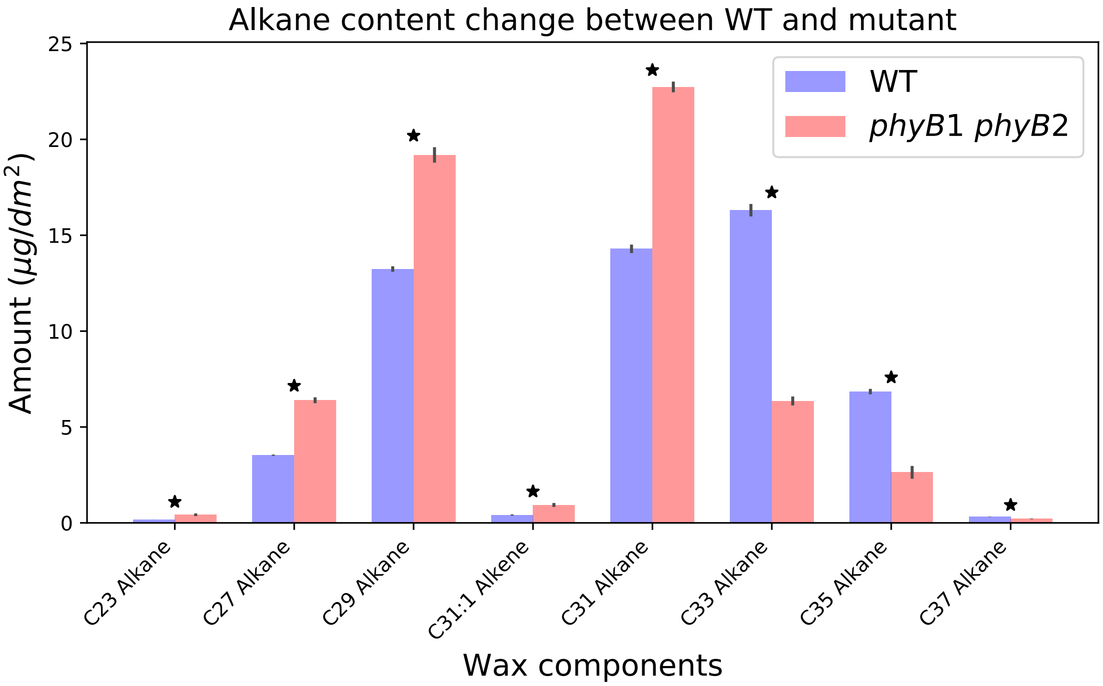


**Fig. S5.** Comparisons of alkanes in cuticular wax profiles in *phyB1 phyB2* double mutants versus nonmutant plants in maize. Asterisks indicate significant differences (< 0.05 FDR) between samples. Error bars represent standard errors.

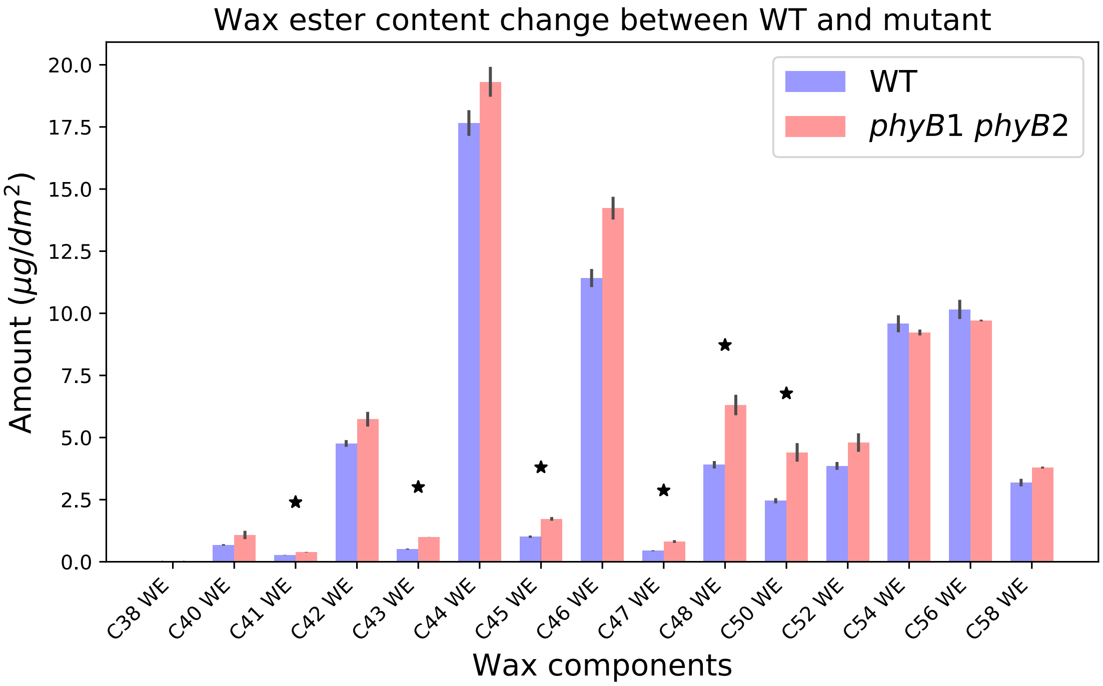


**Fig. S6.** Comparisons of wax esters in cuticular wax profiles in *phyB1 phyB2* double mutants versus nonmutant plants in maize. Asterisks indicate significant differences (< 0.05 FDR) between samples. Error bars represent standard errors.

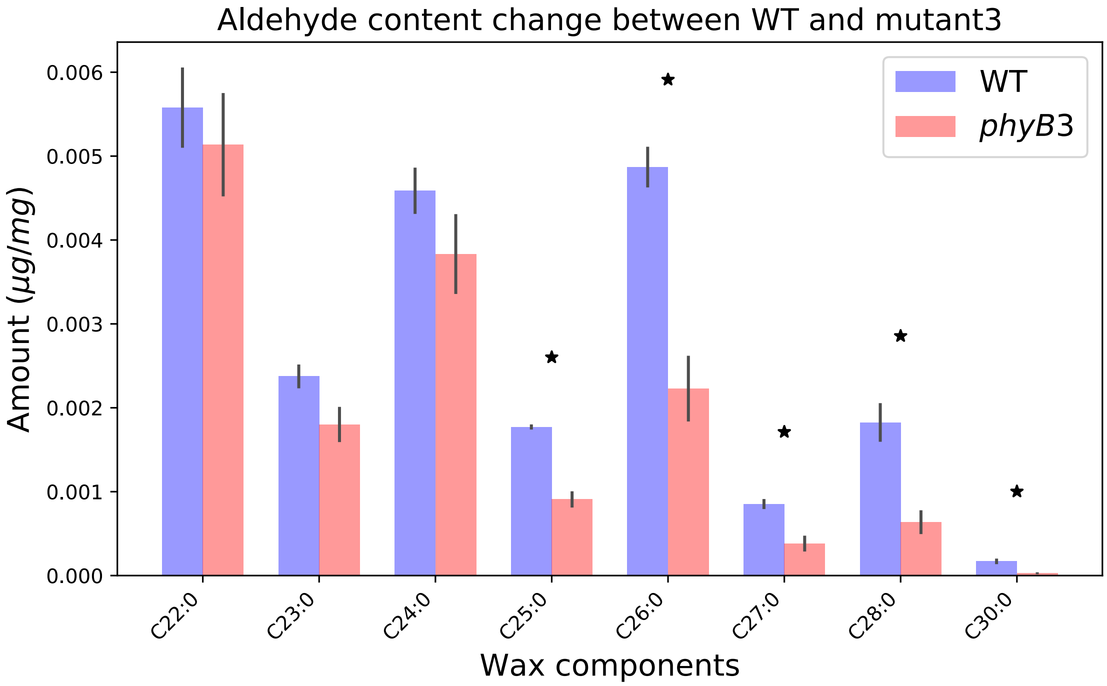


**Fig. S7.** Comparisons of aldehydes in cuticular wax profiles in *phy3* mutants versus nonmutant plants in moss *P. patens.* Asterisks indicate significant differences (< 0.05 FDR) between samples. Error bars represent standard errors.

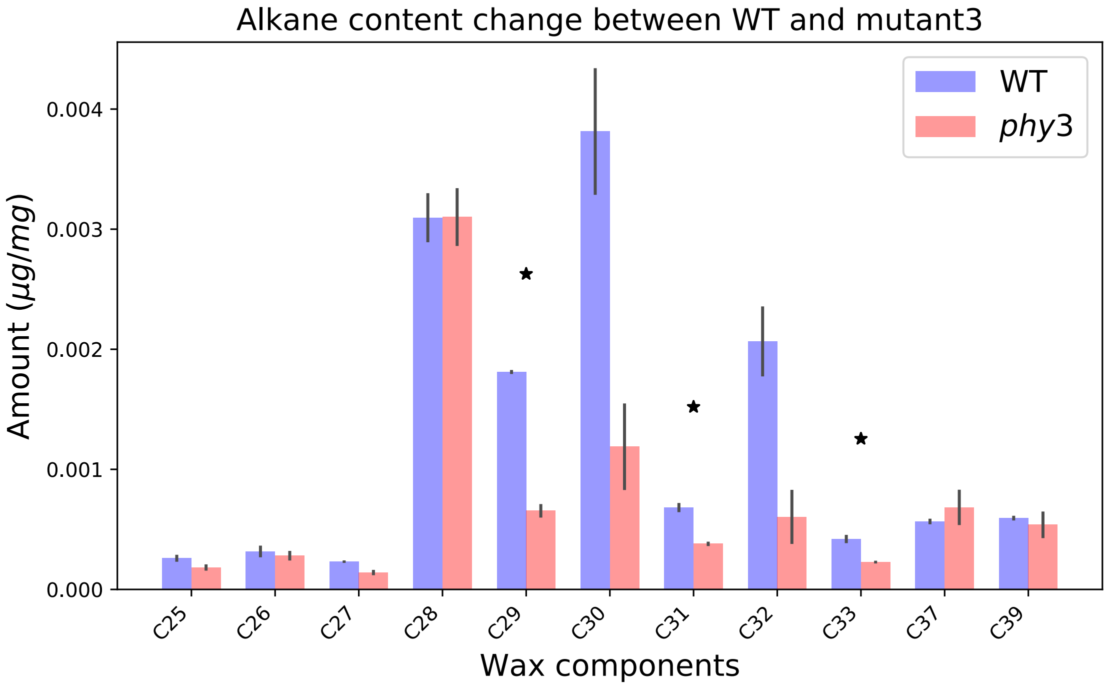


**Fig. S8.** Comparisons of alkanes in cuticular wax profiles in *phy3* mutants versus nonmutant plants in moss *P. patens.* Asterisks indicate significant differences (< 0.05 FDR) between samples. Error bars represent standard errors.

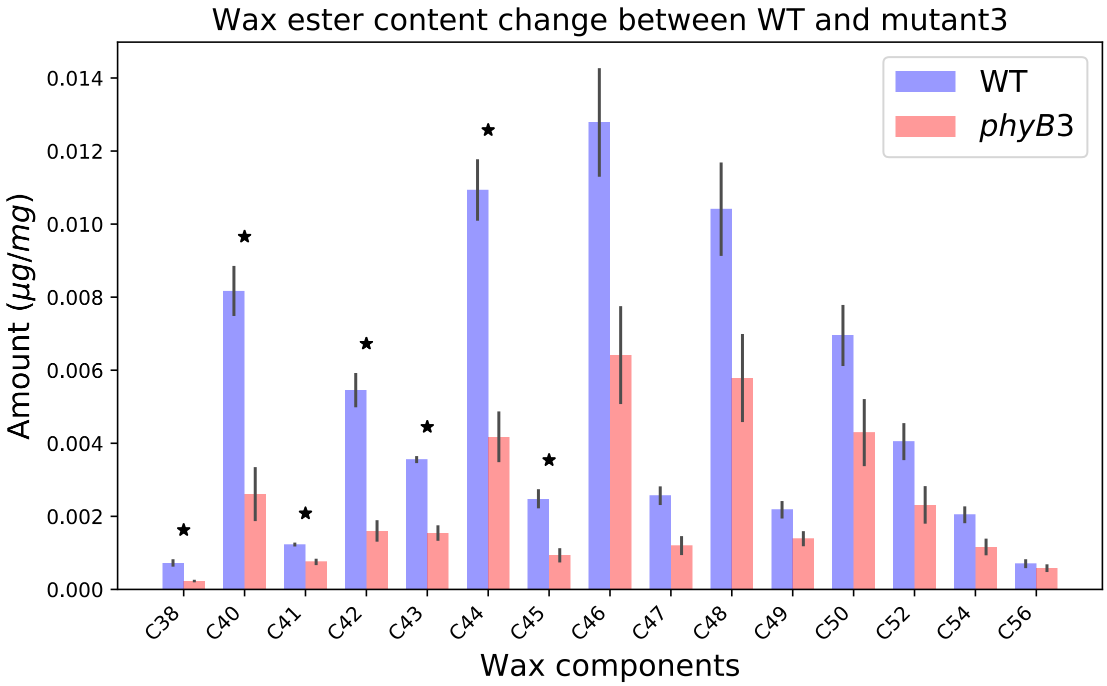


**Fig. S9.** Comparisons of wax esters in cuticular wax profiles in *phy3* mutants versus nonmutant plants in moss *P. patens.* Asterisks indicate significant differences (< 0.05 FDR) between samples. Error bars represent standard errors.
