## Supplementary figures and images for "Network analyses implicate a role for PHYTOCHROME-mediated light signaling in the regulation of cuticle development in plant leaves"

### Supplmental Figure 3

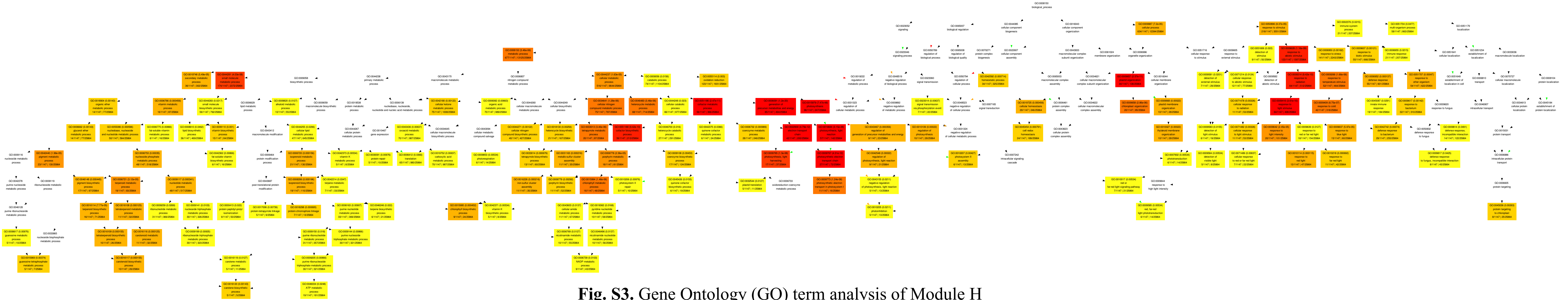

**Fig. S3.** Gene Ontology (GO) term analysis of Module H
